# The alkyl side chain of PACA nanoparticles dictates the impact on cellular stress responses and the mode of particle-induced cell death

**DOI:** 10.1101/304618

**Authors:** Marzena Szwed, Tonje Sønstevold, Anders Øverbye, Nikolai Engedal, Beata Grallert, Ýrr A Mørch, Tore-Geir Iversen, Tore Skotland, Kirsten Sandvig, Maria L Torgersen

**Affiliations:** Department of Molecular Cell Biology, Institute for Cancer Research, Oslo University Hospital, Montebello, N-0379 Oslo, Norway.; Centre for Molecular Medicine Norway (NCMM); Nordic EMBL Partnership; University of Oslo, Norway.; Department of Radiation Biology, Institute for Cancer Research, Oslo University Hospital, Montebello, N-0379 Oslo, Norway.; SINTEF Materials and Chemistry, Sem Sælands vei 2A, 7034 Trondheim, Norway.; Department of Biosciences, Faculty of Mathematics and Natural Sciences, University of Oslo, 0316 Oslo, Norway.

**Keywords:** Poly(alkylcyanoacrylate), nanoparticle, ER stress, oxidative stress, integrated stress response, ferroptosis, Nrf2, ATF4

## Abstract

For optimal exploitation of nanoparticles (NPs) in biomedicine, and to predict nanotoxicity, detailed knowledge on the cellular responses to cell-bound or internalized NPs is imperative. The outcome of NP-cell interaction is dictated by the type and magnitude of the NP insult and the cellular response. Here, we have systematically studied the impact of minor differences in NP composition on cellular stress responses and viability by using highly similar poly(alkylcyanoacrylate) (PACA) particles. Surprisingly, PACA particles differing only in their alkyl side chains; butyl (PBCA), ethylbutyl (PEBCA), or octyl (POCA), respectively, induced different stress responses and modes of cell death in human cell lines. POCA particles induced endoplasmic reticulum stress and apoptosis. In contrast, PBCA and PEBCA particles induced lipid peroxidation by depletion of the main cellular antioxidant glutathione (GSH), in a manner depending on the levels of the GSH precursor cystine, and transcription of the cystine transporter SLC7A11 regulated by ATF4 and Nrf2. Intriguingly, these particles activated the recently discovered cell death mechanism ferroptosis, which constitutes a promising alternative for targeting multidrug-resistant cancer stem-like cells. Of the two, PBCA was the strongest inducer. In summary, our findings highlight the cellular sensitivity to nanoparticle composition and have important implications for the choice of PACA monomer in therapeutical settings.

## Introduction

The fact that many normally ‘inert’ materials become substantially more reactive when downsized to NPs can be beneficial in nanomedicine, but also give rise to nanotoxicity [1]. Intracellular uptake of NPs into lysosomes, or even NP binding to the cell surface, has been shown to activate cellular stress responses, such as oxidative stress or endoplasmic reticulum (ER) stress [2–4], as illustrated in Figure 1A. In NP-mediated drug delivery, cellular stress pathways that are induced as a response to the nanocarrier itself may be beneficial if these stress responses sensitize the target cells. A variety of NPs have been shown to alter the cellular redox balance either by overproduction of reactive oxygen species (ROS) or by depleting the cellular reserve of the main ROS scavenger molecule, reduced glutathione (GSH) [2]. To regain redox homeostasis, an antioxidant response is initiated by activation of transcription factors like Nrf2, HIF1, NFkB, or ATF4, which leads to increased expression of proteins with antioxidant properties [5, 6]. ER stress is characterized by disruption of ER lumen homeostasis, which leads to accumulation of unfolded proteins in the ER. This activates the unfolded protein response (UPR), an adaptive pathway that aims to clear unfolded proteins and restore ER homeostasis [7]. A diversity of NPs has been reported to induce accumulation of ubiquitinated proteins or activate the UPR [3].

**Figure 1:**
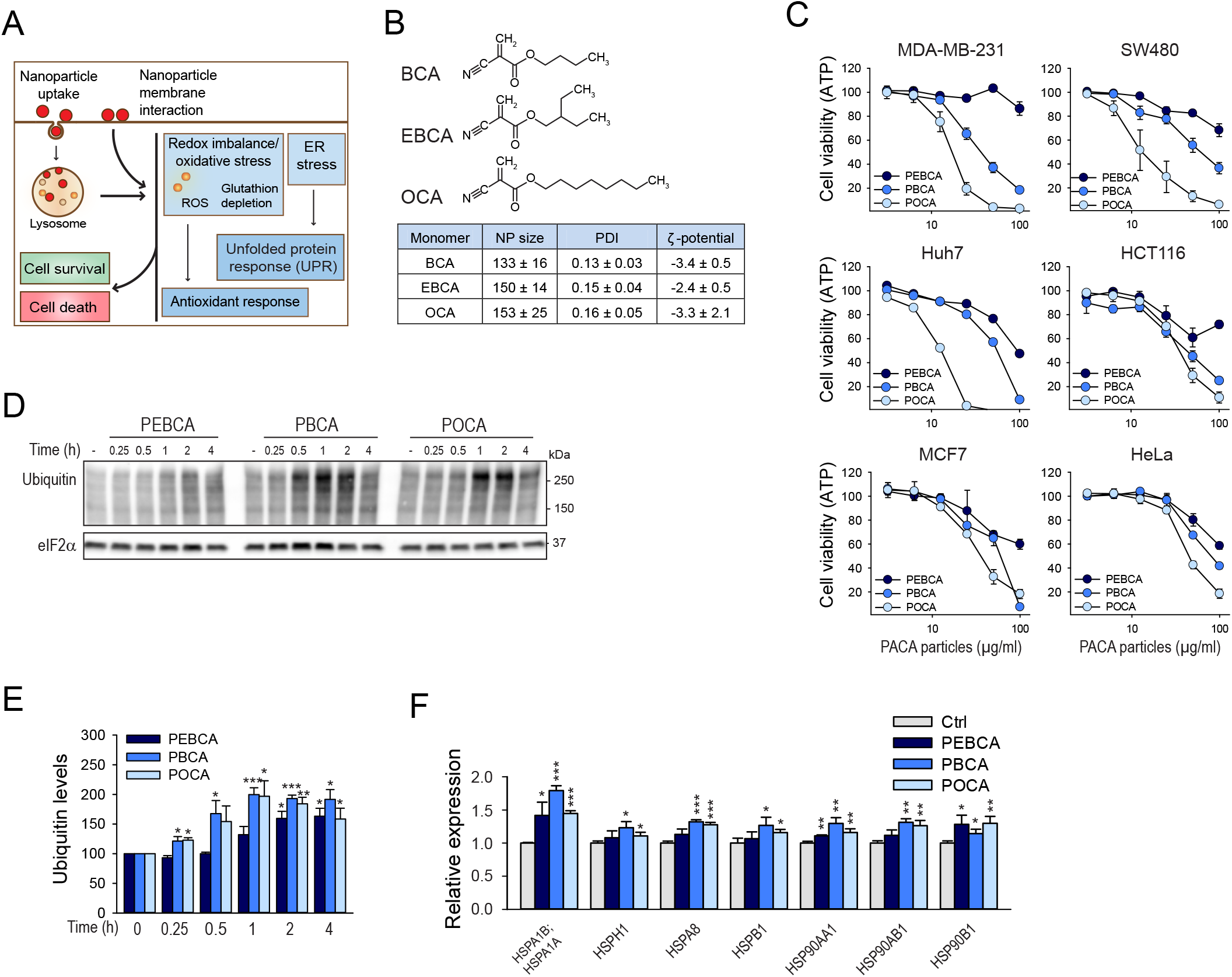
Highly similar PACA particles induce differential cytotoxicity and heat shock response. **A**. Schematic overview of common NP-induced cellular stress responses that contribute to the final outcome of NP-cell interactions. **B**. Molecular structure of the highly similar alkylcyanoacrylate monomers used in this study, and physicochemical characteristics of the resulting NPs. **C**. Cell viability after 24 hours of treatment with PEBCA, PBCA or POCA particles was assessed by determining the cellular ATP levels. 2-fold dilutions of particles were added, from 100 to 3.12 μg/ml. All values were normalized to that of untreated control cells. **D**. MDA-MB-231 cells were treated with PEBCA, PBCA, or POCA particles (25 μg/ml) for the indicated times and cell lysates were prepared for immunoblotting. The blots were probed with the indicated antibodies, and representative blots are shown. **E**. Relative expression of ubiquitin normalized to total eIF2α. **F**. Relative expression of selected heat shock proteins after treatment of HCT116 cells with PEBCA, PBCA or POCA particles (25 μg/ml, 4 hours). The graphs show mean values ± SEM quantified from at least three independent experiments, except the MS data that originate from triplicate samples from one experiment, and the viability data for Huh7 cells that originate from one experiment in triplicate. The asterisks denote the statistical significances compared to untreated control, unless otherwise indicated. *, p< 0.05; **, p<0.01; ***, p< 0.001.

Both the UPR and the antioxidant response are initially pro-survival pathways induced to regain cellular homeostasis. However, if the insult is too severe or prolonged the antioxidant response can no longer protect the cells, and the UPR response is switched to a pro-death pathway [8]. Thus, NP-induced stress pathways are two-faced; they may positively contribute to the intended NP-based therapy in certain settings, but may also prevent an optimal effect in others. Moreover, excessive stress responses may give rise to NP cytotoxicity in normal tissues [1]. Thus, detailed knowledge on the nature and extent of NP-induced cellular stress responses is required to fully exploit the potential of NPs in therapy.

Poly(alkylcyanoacrylate) (PACA), originally used as surgical glue, is a promising drug carrier material [9]. PACA NPs have the ability to cross the blood-brain barrier [10], and are currently in Phase III clinical trials for treatment of advanced hepatocellular carcinoma (ClinicalTrials.gov NCT01655693). Particles are prepared by *in situ* polymerization of alkylcyanoacrylate monomers, and NPs with slightly different chemical compositions can be obtained by using monomers differing only in their alkyl side chains [11, 12]. Here, we have taken advantage of this to study whether minor alterations in NP composition can affect the type or magnitude of cellular stress responses, and whether this may also affect particle cytotoxicity or even the mode of cell death. We have previously shown that particles synthesized from the monomers butyl-, ethylbutyl-, or octyl cyanoacrylate, termed PBCA, PEBCA, and POCA, respectively, exhibit similar physicochemical characteristics, such as size, polydispersity, and surface charge [11, 12]. Intriguingly, we here find that the subtle differences of the alkyl side chains of cyanoacrylate dictate both the type and extent of particle-induced stress responses, and determine the mode of cell death induced by the PACA particles. POCA particles potently induced ER stress and apoptosis. In contrast, PBCA particles induced oxidative stress and ferroptosis, a recently described necrotic cell death pathway, which constitutes a promising alternative cell death pathway in apoptosis-resistant cancer cells, including cancer stem cells [13–15]. The PEBCA particles affected the cells less severely than PBCA and POCA particles, but induced similar phenotypes as PBCA when the availability of the GSH precursor cystine was lowered. Thus, the highly similar PACA particles differentially activated cellular stress, and the final outcome of the PACA treatment depended on the cells’ ability to mount protective stress responses.

## Materials and methods

### Materials

ISRIB, Hoechst 33342, thapsigargin (TG), cycloheximide, H_2_O_2_, buthionine sulfoximine (BSO), reduced glutathion (GSH), N-acetyl cysteine (NAC), liproxstatin, ferrostatin, deferiprone, L-methionine, L-cystine, β-mercaptoethanol, and trichloroacetic acid were all from Sigma. zVAD-fmk was from EMD Millipore Corporation, and LLOMe (SC-285992) was from Santa Cruz.

### Nanoparticle synthesis

PACA NPs were prepared using the mini-emulsion polymerization method as previously described [11]. Briefly, PBCA, PEBCA, or POCA NPs were made by mixing the oil phase, consisting of the monomers butyl-, ethylbutyl-, or octyl cyanoacrylate, a neutral oil, and the fluorescent dye pentamer hydrogen thiophene acetic acid methyl ester (pHTAM), with the aqueous phase consisting of hydrochloric acid and the PEG-surfactants Kolliphor^®^ HS15 and Pluronic^®^ F68. The oil in water mini-emulsion was made using a tip sonifier. The polymerization was carried out overnight and unreacted monomer and surplus of surfactants were removed by extensive dialysis. The size, size distribution and ζ-potential were determined using dynamic and electrophoretic light scattering (Zetasizer Nano ZS, Malvern Instruments) in 0.01 M phosphate buffer, pH 7. The reported NP mean diameter is the Z-average.

### Cells and treatments

The cell lines MDA-MB-231, HCT116, SW480, Huh-7, MCF7, and HeLa were obtained from ATCC and maintained in either RPMI or DMEM containing GlutaMAX^™^ and supplemented with 10% fetal calf serum, 100 U/ml penicillin, and 100 μg/ml streptomycin at 37°C under 5% CO_2_. All cells were seeded one day prior to the experiments. For inhibitor studies, cells were pretreated with the inhibitor at the indicated concentration for 30 minutes before addition of PACA particles. The cellular ATP level (as a surrogate for viability) was measured using the CellTiter-Glo^®^ Luminescent Cell Viability Assay (Promega) and cell death was measured by the CellTox^™^ Green Cytotoxicity Assay (Promega) according to the manufacturer’s procedures. Both luminescence and fluorescence were measured by a Synergy2 plate reader (BioTek). Cystine-free medium was prepared by supplementing DMEM without cystine and methionine (#21013, Gibco) with L-methionine (200 μM), 10% fetal calf serum, 4 mM GlutaMAX^™^ (Gibco), 100 U/ml penicillin, and 100 μg/ml streptomycin. Determination of protein synthesis was performed after 4 hours of treatment with PACA particles, by incubating the cells with leucine-free HEPES-buffered medium complemented with 2 μCi/ml [^3^H]leucine (PerkinElmer) at 37°C for 20 min before proteins were precipitated with 5% (w/v) trichloroacetic acid and washed once with the same solution. Finally, the proteins were dissolved in 0.1 M KOH and radioactively labeled leucine-incorporation was quantified by β-counting with a Tri-Carb 2100TR^®^ Liquid Scintillation Analyzer (Packard Bioscience).

### Immunoblotting

For preparation of total cell lysates treated cells were washed with PBS and lysed directly in 1.1 x Laemmli sample buffer. The lysate was boilt and sonicated briefly to reduce viscosity. Lysates were separated by 4-20% SDS-PAGE and transferred to a PVDF membrane. The membrane was blocked by drying followed by overnight incubation with the indicated primary antibodies in 5% BSA, 35 minutes incubation with HRP-conjugated secondary antibodies and detection with SuperSignal West Dura Extended Duration Substrate (Thermo Scientific) in a ChemiDoc Imaging System (Bio-Rad). The signal intensities were quantified by the Quantity One software (Bio-Rad) and were normalized to the loading control. The following antibodies were used: Ubiquitin (#3936), eIF2α (#5324), PERK (#5683), phospho-eIF2α^Ser51^ (p-eIF2α, #3398), ATF4 (#11815), GCN2 (#3302), PARP (#9542), cleaved caspase-3 (#9661), phospho-p38^Thr180/Tyr182^ (p-p38, #9211), phospho-c-Jun (p-c-Jun, #9164), phospho-p70 S6K^Thr389^ (p-S6K, #9205), p70 S6K (S6K, #9202), and xCT/SLC7A11 (#12691) all from Cell Signaling Technology, and p38 (#612168, Transduction Laboratories), GAPDH (#ab9482, Abcam), XBP1s (#619502, BioLegend), PKR (#ab32052, Abcam), phospho-GCN2 (p-GCN2, #AF7605, R&DSystems) and Actin (#CLT9001, Cedarlane).

### Mass spectrometry analysis

Cells were treated with 25 μg/ml of PBCA, PEBCA or POCA particles and after 4 hours incubation the medium was removed and cells lysed with urea buffer (8 M urea, 2 M thiourea, 50 mM NaCl, 12.5 mM Tris-Glycine, protease and phosphatase inhibitors). To include detached cells, the medium was aspirated and centrifuged at 10,000 x g and resulting pellets combined with cell lysates. Crude cell extracts were sonicated and centrifuged to remove cell debris. Cell lysates were digested with trypsin and obtained peptides subjected to LC/MS/MS separation and identification by an LTQ-Orbitrap XL followed by MaxQuant analysis essentially as described in Hessvik *et al*.[16].

### Determination of reactive oxygen species (ROS) production

Intracellular ROS production was detected by the chloromethyl derivative of the fluorogenic dye 2’,7’-dichlorodihydrofluorescein diacetate (CM-H2DCF-DA, Invitrogen) according to the manufacturer’s procedure. Cells (1.5 × 10^4^ cells/well) were pre-incubated with the dye (10 μM, 45 minutes), rinsed and placed in fresh medium without phenol red in the presence or absence of N-acetyl cysteine (NAC, 5 mM). Subsequently, the cells were treated for 30 minutes with PACA particles, or H_2_O_2_ (100 μM) as a positive control. Fluorescence intensity was measured by a Synergy2 plate reader (BioTek) with excitation and emission wavelengths of 485 nm and 528 nm, respectively.

### Glutathione measurement

The intracellular concentration of reduced glutathione (GSH) was evaluated with o-phthalaldehyde essentially as previously described [17]. The PACA-treated cells (5 × 10^5^) were harvested by Accutase^®^ Cell Detachment Solution (Sigma-Aldrich), washed with PBS, lysed (0.1 M NaCl, 10 mM Na_2_HPO_4_, pH 7.4, 1 mM EDTA, 1% Triton X-100, 60 mM n-octyl β-D-glucopyranoside, and cOmplete EDTA-free Protease Inhibitor Cocktail (Roche)) and precipitated with RQB-TCA solution (20 mM HCl, 5 mM diethylene triamine pentaacetic acid, 10 mM ascorbic acid, 10% trichloroacetic acid) on ice for 30 minutes. Subsequently, the cell lysates were centrifuged (10 minutes, 13000 rpm) and the supernatants were incubated for 30 minutes at room temperature with 50 μl of phosphate buffer (pH 8.0) containing o-phthalaldehyde in methanol (5 mg/ml). The fluorescence intensity of the formed complex was measured by a Synergy2 plate reader (BioTek) with excitation and emission wavelengths of 360 nm and 460 nm, respectively. Inhibition of glutathion synthesis by buthionine sulfoximine (BSO, 100 μM) for 16 hours was used as a positive control for GSH depletion. The GSH content was normalized to the protein content of each lysate, as determined by the BCA method (Pierce).

### Lipid peroxidation detection

Lipid peroxidation was measured using the dye C11-BODIPY(581/591, Thermo Fisher Scientific). Cells treated with PACA NPs for 4 hours were labeled with C11-BODIPY (2.5 μM, 30 minutes). Subsequently, the stained cells were harvested by Accutase^®^ Cell Detachment Solution (Sigma-Aldrich), pelleted, resuspended in PBS and subjected to flow cytometry analysis. The dye was excited using a 488 nm Ar laser and detected with the FL1 (545 nm) detector on an LSR II Flow Cytometer (BD Bioscience). At least 10 000 cells were acquired for each measurement.

### Quantitative real-time RT-PCR

Total RNA was isolated from cells using the RNeasy Plus Mini Kit (QIAGEN), according to the manufacturer’s procedure. 0.8 μg of total RNA was used for cDNA synthesis using the iScript cDNA Synthesis kit (Bio-Rad Laboratories). The real-time PCR analysis was run on a LightCycler 480 Real-Time PCR System using LightCycler 480 SYBR green 1 Master mix (Roche). The cycling conditions were 95°C for 5 minutes, followed by 45 cycles of 95°C 10 s, 60°C 20 s, and 72°C 10 s. The house-keeping gene TBP (TATA box binding protein) served as an internal control and LightCycler 480 Relative Quantification Software (Roche) was used for quantification. The following QuantiTect^®^Primer Assays were used: Hs_EIF2AK4 (QT01036350), Hs_EIF2AK3 (QT00066003), Hs_EIF2AK2 (QT00022960), Hs_EIF2AK1 (QT01018920), Hs_ATF4 (QT00074466), Hs_TBP (QT00000721), Hs_SLC7A11 (QT00002674), Hs_GCLM (QT00038710), Hs_FTH1 (QT00072681), and Hs_HMOX (QT00092645).

### Transfection of cells with siRNAs

Small interfering RNAs (siRNAs) were introduced into MDA-MB-231 cells by reverse transfection. The siRNAs were diluted in RPMI and mixed with Lipofectamine RNAiMax (Invitrogen) before addition to trypsinized cells at a final siRNA-concentration of 10 nM. 3 × 10^5^ cells were seeded in 6-well culture plates, and the culture medium was exchanged the next day. The experimental treatments were initiated after 48 h of transfection. The following siRNAs were used (Silencer Select siRNAs from Ambion): Silencer^®^ Select Negative Control #1 (4390843), siPERK-1 (ID 18103), siPERK-2 (ID 18101), siATF4-1 (ID 1702), siATF4-2 (ID 1704), siHRI-1 (ID 25822), siHRI-2 (ID 25823), siNrf2-1 (ID 9492), siNrf2-2 (ID 9493), siSLC7A11-1 (ID 24289), siSLC7A11-2 (ID 24291), siPKR-1 (ID 11185), siPKR-2 (ID 11186), or ON-TARGETplus Non-targeting siRNA #1(D-001810-01-05), ON-TARGET plus siGCN2-1 (J-005314-07-0010) and siGCN2-2 (J-005314-06-0010) from Dharmacon. In the quadruple knockdown of ISR kinases 10 nM of each of the following siRNAs were included: siPERK-1, siGCN2-2, siPKR-1, siHRI-1. This was compared to 40 nM of negative control siRNA. In the double knockdown of ATF4 and Nrf2, 10 nM of each siRNA (siATF4-1 and siNrf2-2) was mixed and compared to 20 nM of negative control siRNA.

### Microscopy

Phase contrast images were acquired by an Eclipse TS100 microscope (Nikon) equipped with a 20x objective and a Digital Sight camera (Nikon). Cell death was visualized by the fluoregenic dye CellTox^™^ Green Cytotoxicity reagent (Promega) after 24 hours treatment of the cells with PACA particles. Merged fluorescent and phase contrast wide-field images were acquired by the EVOS FL Cell Imaging System (Thermo Fisher Scientific). Hoechst-stained nuclei were imaged using a Zeiss LSM780 laser scanning confocal microscope (Carl Zeiss MicroImaging) equipped with a Laser diode 405-30 CW (405 nm). The objective used was a Zeiss Plan-Apochromat 63×/1.40 Oil DIC M27. Images were acquired using the ZEN 2010 software (Carl Zeiss MicroImaging).

For live cell microscopy cells seeded on glass bottom dishes (MatTek) were imaged on a Deltavision microscope (Applied Precision) equipped with Elite TruLight Illumination System, a CoolSNAP HQ2 camera, and a 60× Plan-Apochromat (1.42 NA) objective. The microscope stage was kept at 37°C under 5% CO_2_ by an incubation chamber. For detection of caspase-3/7 activity the PACA particles was added to a 4-chamber MatTek dish in the presence of the fluorogenic dye CellEvent^™^ Caspase 3/7 Green Detection Reagent (Invitrogen), and then time-lapse images (5 z-sections 2.5 μm apart) were acquired every 10 minutes over a total time period of 18 hours. For detection of galectin-3 puncta HeLa cells stably expressing mCherry-Galectin-3 was grown in MatTek dishes, and then a 2x concentrated solution of LLOMe or PACA particles (as indicated in the figure) was injected into the dish at t=5 minutes. Time-lapse images (5 z-sections 2.5 μm apart) were acquired every 1 minute over a total time period of 1 hour for LLOMe treatment, and every 5 minutes over a total time period of 18 hours for PACA-treatment. The images were deconvolved and z-projected using the softWoRx software (Applied Precision).

### Statistical analysis

Mean values ± standard error of the mean (SEM) were calculated for each condition. The statistical significance of the differences was determined by two-tailed unpaired Student’s t-test, with equal or unequal variances, as appropriate; *, p<0.05; **, p<0.01; ***, p<0.001.

## Results

### The PACA particles exert differential cytotoxicity and induction of heat shock proteins

PACA particles with only minor differences in their cyanoacrylate side chains (Figure 1B) were produced by *in situ* polymerization of the alkylcyanoacrylate monomers butyl-(PBCA), ethylbutyl-(PEBCA), or octyl-cyanoacrylate (POCA) as previously described [11, 12]. The resulting particles exhibited similar mean sizes, in the range of 133-153 nm, with a relatively narrow size distribution. All three NPs were slightly negatively charged with a zeta-potential of approximately −3 mV (Figure 1B). When particle cytotoxicity was compared across a panel of cancer cell lines (MDA-MB-231, SW480, Huh7, HCT116, MCF7, and HeLa) the POCA particles consistently exerted the highest cytotoxicity, whereas PEBCA particles were the least toxic (Figure 1C). This is in agreement with our previous screening study with a set of 12 other cancer cell lines [11]. Importantly, we have found that this pattern of cytotoxicity is not caused by gross differences in cellular uptake of the particles [11, 18]. To assess whether the differential PACA cytotoxicity is caused by particle-specific induction of cellular stress responses, we focused on the triple negative breast cancer cell line MDA-MB-231 that displayed the largest diversity in particle cytotoxicity (Figure 1C).

We first assessed whether the PACA particles would induce accumulation of ubiquitinated proteins, a hallmark of altered cellular homeostasis [7]. When cells were treated with 25 μg/ml (an intermediary concentration with respect to cytotoxicity at 24 hours) of each variant of PACA NPs, a buildup of ubiquitinated proteins was observed within the first 4 hours of treatment in all cell lines tested (Figure 1D-E, Supplementary Figure 1A). The induction of ubiquitin-like modifications was verified by mass spectrometry analyses (Supplementary Figure 1B). The PACA particles also induced increased expression of a wide variety of heat shock family proteins, a hallmark of cellular stress response (Figure 1F). Overall, the PBCA and POCA particles exerted the most potent effect.

### The PACA particles differentially induce the unfolded protein response

To identify the nature of the PACA-induced cellular stress, we first assessed activation of ER stress and the UPR using the PERK pathway as readout (Figure 2A). During ER stress, accumulation of unfolded proteins in the ER lumen activates PERK, and subsequently, global protein synthesis is shut down via phosphorylation, and thus inactivation, of eukaryotic translation initiation factor 2 (eIF2α) [7]. This allows for preferential translation of UPR-dependent genes, such as the activating transcriptional factor 4 (ATF4), which initiates transcription of genes encoding ER chaperones, amino acid transporters, and antioxidant proteins that contribute to restoration of cellular homeostasis [7]. Intriguingly, the three variants of PACA NPs induced a highly different UPR activation pattern (Figure 2B-D). At 25 μg/ml the PEBCA particles modestly elevated phospho-eIF2α levels in most cell lines (Figure 2B-C, Supplementary Figure 2A). Higher PEBCA concentrations induced PERK phosphorylation, as detected by an upward shift in PERK migration (Supplementary Figure 2B). In contrast, 25 μg/ml of POCA particles induced phosphorylation of PERK in all cell lines (Figure 2B, Supplementary Figure 2A-B). Surprisingly, 25 μg/ml of PBCA particles induced potent phosphorylation of eIF2α and accumulation of ATF4 without a clear upstream PERK activation (Figure 2B-D, Supplementary Figure 2A), and a less marked shift than observed with PEBCA and POCA at elevated concentrations (Supplementary Figure 2B). This might suggest that eIF2α is predominantly phosphorylated by an alternative kinase in PBCA-treated cells. To assess whether the particles activate the IRE1 ER stress pathway, we probed for the IRE1 downstream target XBP1s [7]. An accumulation of XBP1s was detected upon treatment with PEBCA particles and low POCA concentrations, whereas PBCA particles did not induce XBP1s (Supplementary Figure 2B). Together, this suggests that ER stress is induced by the POCA particles, and also by PEBCA particles at high concentrations, but not by PBCA particles.

**Figure 2.**
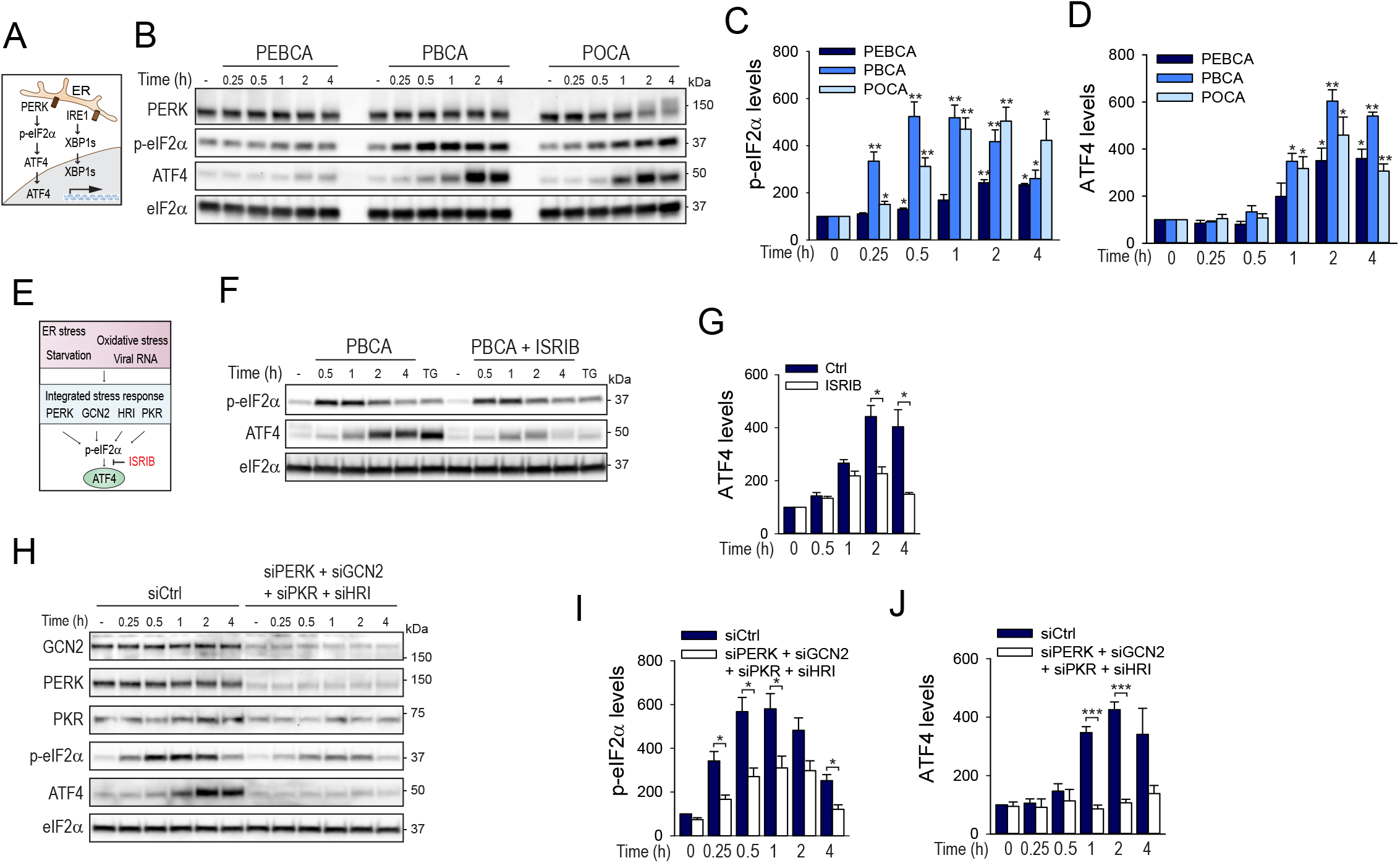
PBCA-induced ATF4 accumulation is dependent on the integrated stress response. **A**. Schematic overview of the PERK - and IRE1 ER stress pathways. **B**. MDA-MB-231 cells were treated with PEBCA, PBCA, or POCA particles (25 μg/ml) for the indicated times and cell lysates were prepared for immunoblotting. The blots were probed with the indicated antibodies, and representative blots are shown. The relative levels of p-eIF2α (C), and ATF4 (D) were normalized to total eIF2α. **E**. Schematic overview of the integrated stress response consisting of the four eIF2α kinases PERK, GCN2, HRI and PKR that sense cellular stress, such as ER stress, starvation, oxidative stress, or viral RNA. The downstream effects of phospho-eIF2α are inhibited by ISRIB. **F**. MDA-MB-231 cells were treated with PBCA particles (25 μg/ml) in the absence or presence of ISRIB (100 nM) and cell lysates were prepared for immunoblotting. Thapsigargin (TG, 100 nM) was used as a positive control. The relative levels of ATF4 were normalized to total eIF2α (G). MDA-MB-231 cells were transfected with non-targeting control siRNA or a mixture of siRNAs targeting the four ISR kinases. After 48 hours the depleted cells were treated with PBCA particles (25 μg/ml) for the indicated times and cell lysates were prepared for immunoblotting (H). The relative levels of p-eIF2α (I) and ATF4 (J) were normalized to total eIF2α. All bars show mean values ± SEM quantified from at least three independent experiments. The asterisks denote the statistical significances compared to untreated control, unless otherwise indicated. *, p< 0.05; **, p<0.01; ***, p< 0.001.

### PBCA-induced ATF4 accumulation is mediated by the integrated stress response

Accumulation of the transcription factor ATF4 can not only be observed as a response to ER stress and PERK activation, but also downstream of three other eIF2α kinases, GCN2, PKR and HRI, through the so-called *integrated stress response* (ISR) [19], as illustrated in Figure 2E. These kinases sense various internal or external stressors, including amino acid starvation, viral infection, proteasomal inhibition, and redox imbalance [19]. In order to see whether the PBCA-induced ATF4 accumulation is mediated via the ISR, we employed the recently developed ISR inhibitor ISRIB, which blocks all downstream effects of phosphorylated eIF2α [20] (Figure 2E). Indeed, the PBCA-induced accumulation of ATF4 was inhibited by ISRIB (Figure 2F-G). Thapsigargin was used as a positive control for ISR-induction [21]. In an attempt to identify the specific ISR kinase responsible for the PBCA-induced stress response, the four kinases were depleted separately by siRNAs. Single knockdown of each ISR kinase had little (HRI and GCN2) or no (PERK and PKR) effect on phosphorylation of eIF2α or accumulation of ATF4 (Supplementary Figure 3A-D). However, simultaneous depletion of all four ISR kinases (Supplementary Figure 3E-F) totally abolished PBCA-induced accumulation of ATF4 and strongly reduced the phosphorylation of eIF2α (Figure 2H-J). Together, these data indicate that the PBCA particles activate the ISR.

### PBCA particles alter cellular redox balance, and both particle-induced stress responses and cytotoxicity are prevented by reduced glutathione

As oxidative stress has been implicated in activation of the ISR via GCN2, PKR, and HRI [19], we asked whether PACA particles alter the cellular redox balance. Cellular ROS levels were significantly increased by both PEBCA and PBCA, whereas only a slight tendency of an increase, that did not reach statistical significance, was observed with POCA (Figure 3A). We chose PBCA particles to analyze the nature of this effect further. The main cellular antioxidant, reduced GSH, was partially depleted upon treatment with PBCA (Figure 3B). Thus, PBCA clearly induced redox imbalance, and this seemed to trigger the downstream stress responses, as treatment with excess GSH prevented PBCA-induced accumulation of ubiquitinated proteins, phosphorylation of eIF2α, and accumulation of ATF4 (Figure 3C-F). Also the cytotoxicity of PBCA particles was fully reversed by either excess GSH or NAC, a precursor of GSH [22] (Figure 3G). In contrast, POCA cytotoxicity was not prevented by antioxidant treatment (Figure 3G), pointing to a fundamental difference in the cytotoxic action of these particles.

**Figure 3.**
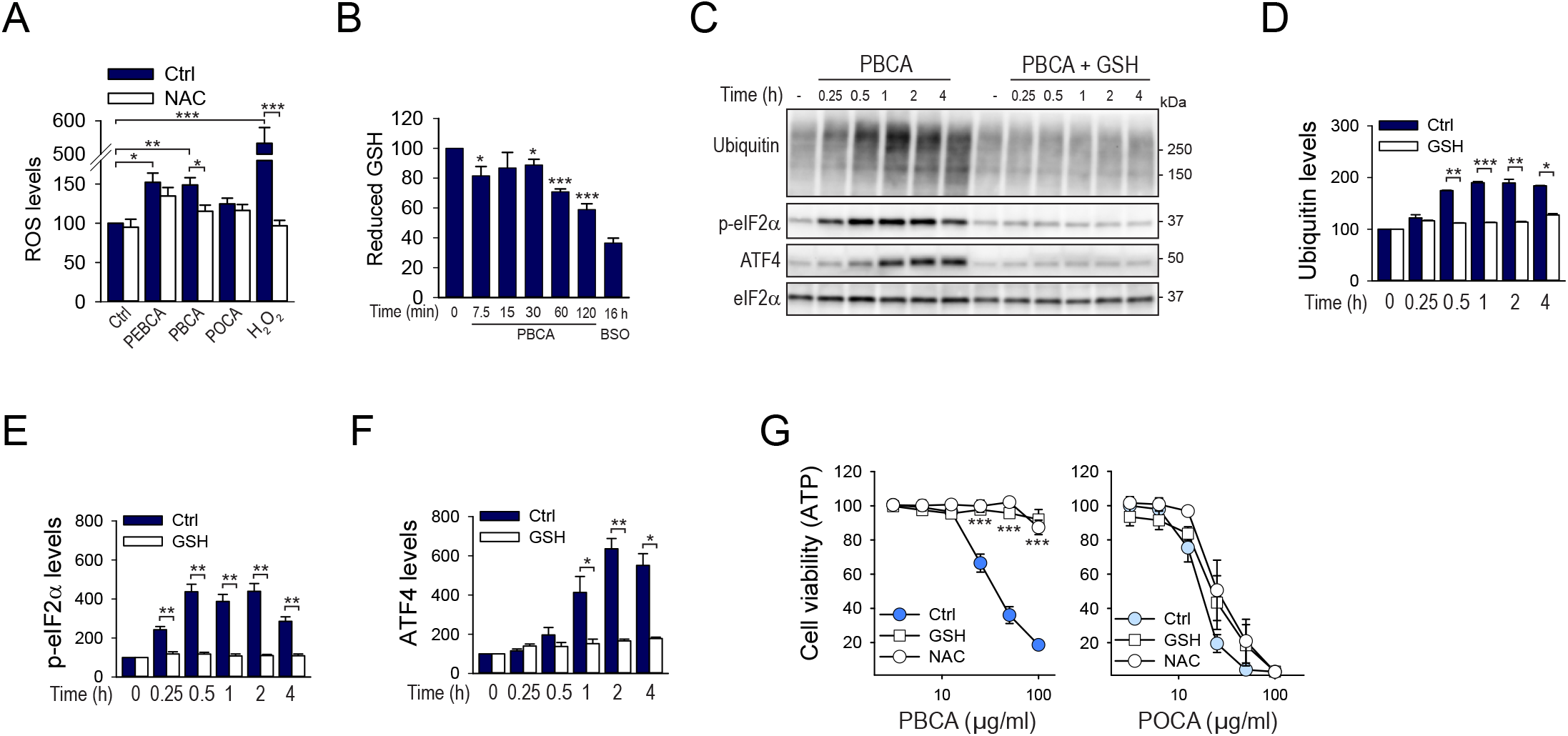
PBCA-induced stress responses and cytotoxicity are mediated by GSH depletion. **A**. The cellular ROS level was assessed by H_2_DCFDA fluorescence after treatment of MDA-MB-231 cells with PEBCA, PBCA or POCA particles (25 μg/ml) for 30 minutes in the absence or presence of N-acetyl cysteine (NAC, 5 mM). H_2_O_2_ (100 μM) was used as positive control. **B**. The levels of reduced glutathione (GSH) was assessed in cells treated with PBCA particles (25 μg/ml) for the indicated times. Inhibition of glutathion synthesis by buthionine sulfoximine (BSO, 100 μM) for 16 hours was used as a positive control for GSH depletion. **C**. MDA-MB-231 cells were treated with PBCA (25 μg/ml) in the absence or presence of reduced glutathione (GSH, 10 mM) for the indicated times and lysates were prepared for immunoblotting. The blots were probed with the indicated antibodies, and representative blots are shown. The relative levels of ubiquitin (D), p-eIF2α (E), and ATF4 (F) were normalized to total eIF2α. **G**. Cell viability assessed by ATP levels of MDA-MB-231 cells treated with PBCA or POCA particles for 24 hours in the absence or presence of either GSH (10 mM) or NAC (5 mM). 2-fold dilutions of particles were added, from 100 to 3.12 μg/ml. All graphs show mean values ± SEM quantified from at least three independent experiments, except for BSO where the error bars represent deviation from the mean of two independent experiments. The asterisks denote the statistical significances compared to untreated control, unless otherwise indicated. *, p< 0.05; **, p<0.01; ***, p< 0.001.

### POCA particles selectively induce apoptotic cell death

Intrigued by these findings, we aimed to identify the mode of cell death induced by the three PACA particle variants. Strikingly, only POCA particles induced an apoptotic phenotype, identified by membrane blebbing, apoptotic bodies and nuclear fragmentation already after 4 hours of treatment (Figure 4A-B). Also other hallmarks of apoptosis, such as cleavage of caspase-3 and PARP, were detected selectively upon treatment with POCA particles in multiple cell lines (Figure 4C, Supplementary Figure 4). PBCA and PEBCA particles instead led to cell rounding and eventual cell swelling and rupture, in the absence of caspase activation, as detected by the fluorogenic CellEvent^™^ caspase 3/7 substrate (Figure 4D, Supplementary videos 1-3). Finally, treatment with the pan-caspase inhibitor z-VAD significantly counteracted POCA cytotoxicity, whereas PBCA cytotoxicity was unchanged (Figure 4E), demonstrating that the highly similar PACA particles induce different modes of cell death.

**Figure 4.**
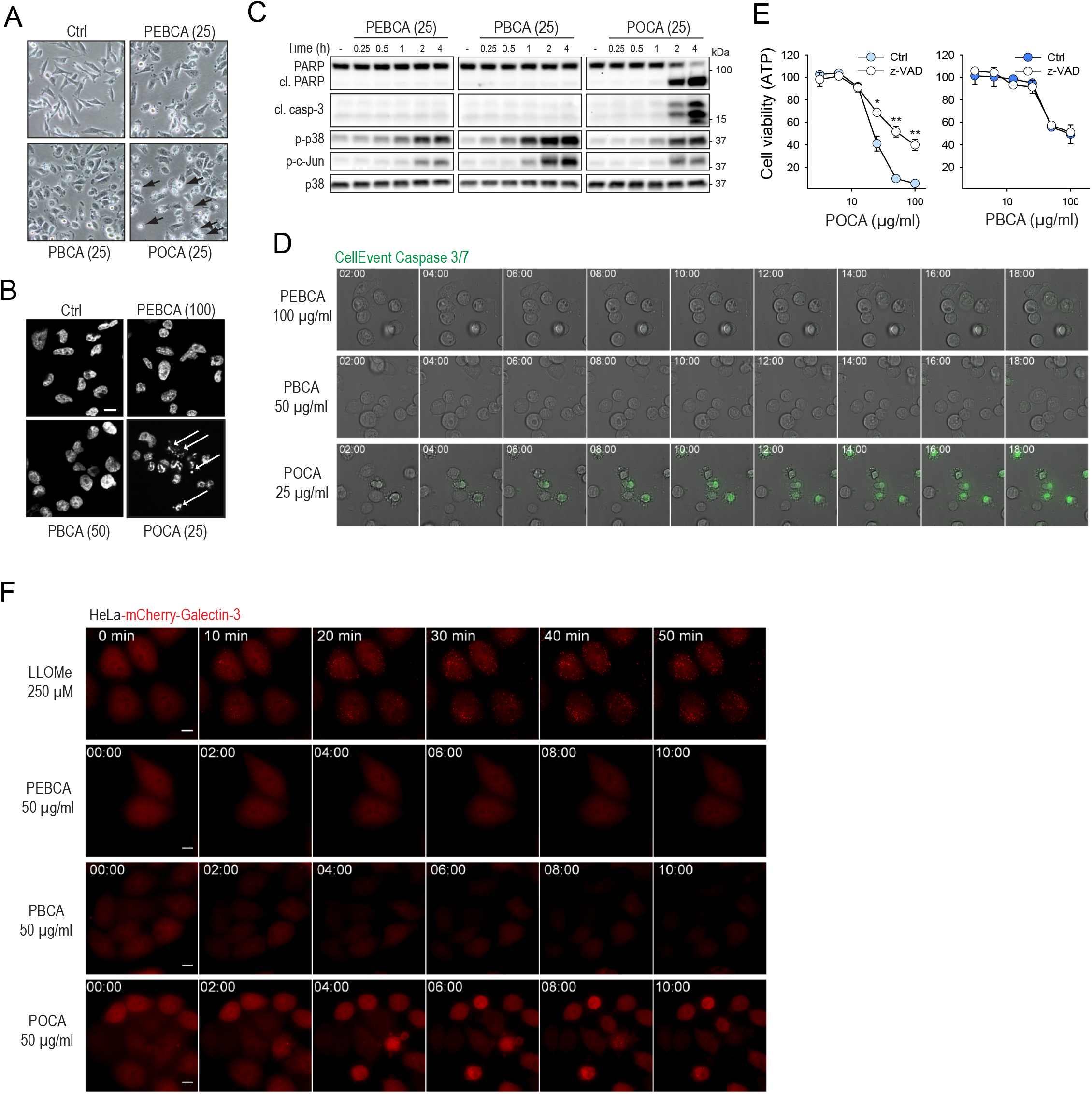
POCA particles selectively induce apoptotic cell death. **A**. MDA-MB-231 cells were treated with the indicated concentrations (μg/ml) of PEBCA, PBCA or POCA particles for 4 hours before morphological assessment (A) or Hoechst staining of the nuclei (B). The arrows indicate apoptotic cells. **C**. MDA-MB-231 cells were treated with PEBCA, PBCA, or POCA particles (25 μg/ml) for the indicated times and cell lysates were prepared for immunoblotting. The blots were probed with the indicated antibodies, and apoptosis is indicated by cleaved PARP (cl. PARP) and cleaved caspase-3 (cl. casp-3). **D**. Caspase-3/7 activity was assessed in MDA-MB-231 cells treated with the indicated concentrations of PEBCA, PBCA or POCA particles for up to 18 hours by employing the fluorogenic substrate CellEvent^™^ Caspase 3/7. **E**. Cell viability assessed by ATP levels of MDA-MB-231 cells treated with PBCA or POCA particles with or without the pan-caspase inhibitor z-VAD (20 μM) for 24 hours. 2-fold dilutions of particles were added, from 100 to 3.12 μg/ml. The mean values of three independent experiments are shown ± SEM. The asterisks denote the statistical significances compared to untreated control *, p< 0.05; **, p<0.01. **F**. Lysosomal membrane permeabilization was assessed in HeLa-mCherry-Galectin-3 cells treated with PEBCA, PBCA or POCA particles (50 μg/ml) for up to 10 hours. LLOMe (250 μM) was used as a positive control. Scale bars; 10 μm.

As permeabilization of the lysosomal membrane has been observed upon treatment with certain NPs [23], it was of interest to see whether the PACA particles induce lysosomal damage. To this end we employed HeLa cells stably expressing mCherry-Galectin-3, a commonly used system for detection of lysosomal permeabilization [24]. As expected, the positive control L-leucyl-L-leucine methyl ester (LLOMe), a lysosomotropic detergent-like substance [24], rapidly induced numerous mCherry-Galectin-3 puncta (Figure 4F). In contrast, the PACA particles, even at high concentrations and prolonged treatment, did not induce evident galectin-3 staining stronger than the background level (Figure 4F, Supplementary videos 4-7). The only exception was galectin-3 puncta appearing in POCA-treated cells several hours after initiation of membrane blebbing, likely as a secondary event of apoptosis. Thus, it seems unlikely that lysosomal membrane permeabilization is the initial event responsible for PACA-induced cell death.

### PBCA and PEBCA particles induce ferroptosis depending on the cellular cystine levels

We next asked what type of non-apoptotic cell death may be induced by PBCA. In recent years, the term necrosis has been subdivided into a plethora of differentially regulated necrotic cell death pathways [25]. A recently discovered subtype of regulated necrosis, called ferroptosis, is associated with accumulation of ROS in an iron-dependent manner, loss of GSH, and excessive lipid peroxidation [13, 14]. As PBCA cytotoxicity seemed to depend on redox imbalance (Figure 3G), we assessed its cytotoxicity in the presence of the ferroptosis inhibitors ferrostatin and liproxstatin, which are lipophilic, small-molecule antioxidants [26]. To selectively label dead cells, we employed the CellTox^™^ Green cytotoxicity assay, in which only cells with loss of plasma membrane integrity are stained. Remarkably, PBCA cytotoxicity was almost totally reversed by the ferroptosis inhibitors (Figure 5A-B). Also the iron chelator deferiprone (DFP) abolished PBCA cytotoxicity, further implicating ferroptosis, whereas the pan-caspase inhibitor z-VAD had no significant effect. As ferroptosis is characterized by lipid peroxidation, we measured the extent of peroxidated lipids by employing the probe BODIPY-C11. Indeed, treatment with PBCA particles induced lipid peroxidation in a liproxstatin-sensitive manner (Figure 5C).

**Figure 5.**
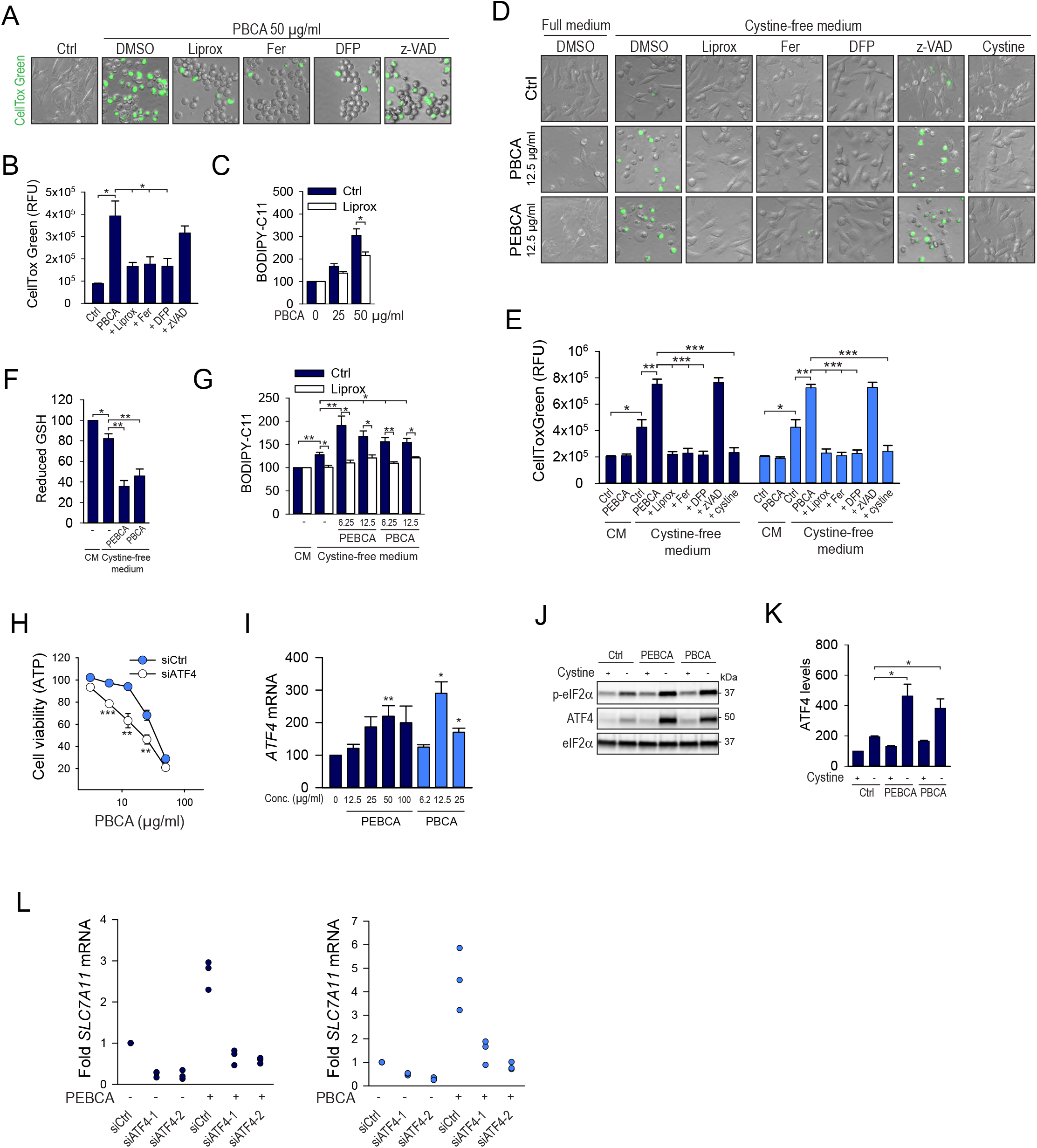
PBCA and PEBCA particles induce ferroptosis in a cystine-dependent manner. **A-B**. Cytotoxicity assessed by CellTox Green staining of MDA-MB-231 cells treated with PBCA particles (50 μg/ml, 24 hours) in the presence or absence of liproxstatin (Liprox, 1 μM), ferrostatin (Fer, 2 μM), deferiprone (DFP, 100 μM), or z-VAD (20 μM). **C**. Flow cytometric determination of BODIPY-C11 staining of MDA-MB-231 cells after 4 hours treatment with the indicated concentrations (μg/ml) of PBCA in the absence or presence of liproxstatin (2 μM). D-E. Cytotoxicity assessed by CellTox Green staining of MDA-MB-231 cells treated with PBCA or PEBCA particles (12.5 μg/ml, 24 hours) in full medium or in cystine-free medium in the presence or absence of liproxstatin (Liprox, 1 μM), ferrostatin (Fer, 2 μM), deferiprone (DFP, 100 μM), z-VAD (20 μM), or cystine (200 μM). **F**. Cellular glutathione levels were determined in cells treated with PEBCA or PBCA particles (12.5 μg/ml, 4 h) in cystine-free medium. **G**. Flow cytometric determination of BODIPY-C11 staining of MDA-MB-231 cells after 4 hours treatment with the indicated concentrations (μg/ml) of PEBCA or PBCA in cystine-free medium in the absence or presence of liproxstatin (1 μM). **H**. Cell viability assessed as ATP levels of MDA-MB-231 cells depleted for ATF4 for 48 hours and then treated with PBCA particles for 24 hours. 2-fold dilutions of particles were added, from 50 to 3.12 μg/ml. The data were normalized to the respective non-particle-treated control for each siRNA. Depletion of ATF4 in itself did not reduce cell viability. **I**. Transcription of *ATF4* mRNA in cells treated with the indicated concentrations of PEBCA or PBCA particles for 4 hours. **J-K**. Cells were treated with PEBCA or PBCA particles (12.5 μg/ml) for 4 hours in full - or cystine-free medium and cell lysates were prepared for immunoblotting. The relative level of ATF4 was normalized to elF2α. **L**. Transcription of *SLC7A11* mRNA in cells transfected with a non-targeting control siRNA (siCtrl) or by two independent siRNAs targeting ATF4.48 hours later the cells were treated with PEBCA (50 μg/ml) or PBCA (12.5 μg/ml) for 4 hours. All graphs show mean values ± SEM quantified from at least three independent experiments. The asterisks denote the statistical significances compared to untreated control, unless otherwise indicated. *, p< 0.05; **, p<0.01; ***, p< 0.001.

The cellular import of cystine is identified as one of the key regulators of ferroptosis [14]. Imported cystine is converted to cysteine, which is the rate-limiting substrate in GSH synthesis. Thus, cystine availability is an important determinant of cellular redox capacity. We thus asked how cystine availability affects PACA cytotoxicity. Strikingly, when MDA-MB-231 cells were incubated in medium lacking only cystine, potent cell death was induced by significantly lower PBCA concentrations than those required to induce cytotoxicity in full medium (Figure 5D-E). The cystine starvation-induced cell death, and importantly also the cell death induced by PBCA particles in cystine-free medium, was fully inhibited by the ferroptosis inhibitors ferrostatin, liproxstatin, or DFP, or by re-addition of cystine (Figure 5D-E), whereas z-VAD had no effect. Surprisingly, in cystine-free medium even the relatively non-cytotoxic PEBCA particles exerted potent ferroptosis-dependent cytotoxicity (Figure 5D-E). To test whether the increased vulnerability towards these two PACA particles was caused by a lower antioxidant capacity in cystine-starved cells, the cellular GSH levels were measured. Indeed, four hours of cystine starvation was sufficient to partially reduce the level of GSH (Figure 5F), and low concentrations of PEBCA and PBCA particles (12.5 μg/ml) further reduced the GSH levels (Figure 5F). Concomitant with loss of GSH, low concentrations of PEBCA and PBCA significantly induced lipid ROS generation in cells starved for cystine (Figure 5G). Taken together, both PEBCA and PBCA seem to have the ability to induce ferroptosis in MDA-MB-231 cells, depending on cystine availability and the resulting cellular antioxidant capacity.

### PEBCA - and PBCA-induced ferroptotic cell death depends on ATF4 and Nrf2

As treatment with PBCA particles potently induced accumulation of ATF4 in a GSH - dependent manner (Figure 3C), we asked whether ATF4 contributes to cell viability in PBCA-treated cells. Indeed, this seemed to be the case. Depletion of ATF4 in MDA-MB-231 cells further potentiated PBCA cytotoxicity at low particle concentrations (up to 25 μg/ml) (Figure 5H, Supplementary Figure 5A), indicating that ATF4 is activated as a pro-survival response. ATF4 was induced at the transcriptional level by low PBCA concentrations, reaching maximum levels at 12.5 μg/ml (Figure 5I). In line with the lower cytotoxicity of PEBCA particles, *ATF4* mRNA was maximally induced at higher PEBCA concentrations (50 μg/ml) (Figure 5I). Moreover, the importance of cystine availability in PBCA/PEBCA - induced stress responses was demonstrated by the significantly higher levels of ATF4 induced under cystine starvation conditions even at low particle concentrations (Figure 5J-K). Importantly, both PEBCA and PBCA particles induced partly ATF4-dependent transcription of *SLC7A11* (Figure 5L), the light-chain subunit of the cystine-glutamate antiporter Xc-, which is responsible for cystine import and regarded as a guardian of the antioxidant defense pathway [14].

As PEBCA and PBCA particles exert comparable cytotoxicity upon cystine starvation (Figure 5E), whereas the particle cytotoxicity is highly different in complete cell medium (Figure 1C), the question remains whether the PEBCA particles have the ability to activate additional cytoprotective pathways. Nrf2 is a master transcriptional activator of cytoprotective genes in response to electrophiles and oxidants [27]. Normally, Nrf2 levels are kept low by ubiquitination via the E3 ubiquitin ligase adaptor Keap1, but the presence of cytotoxic substances blocks the interaction to Keap1 and thus prevents Nrf2 degradation [6]. Interestingly, PEBCA particles more potently activated Nrf2 than PBCA particles; whereas the maximum PBCA-induced Nrf2 activation was reached already at 25 μg/ml and then decreased at higher concentrations, Nrf2 was accumulated in a dose-dependent manner by PEBCA particles (Figure 6A-B), resembling the transcriptional activation of *ATF4* (Figure 5I). This reduced ability to activate cell protective responses at elevated PBCA concentrations may contribute to the differential cytotoxicity of the two particles.

**Figure 6.**
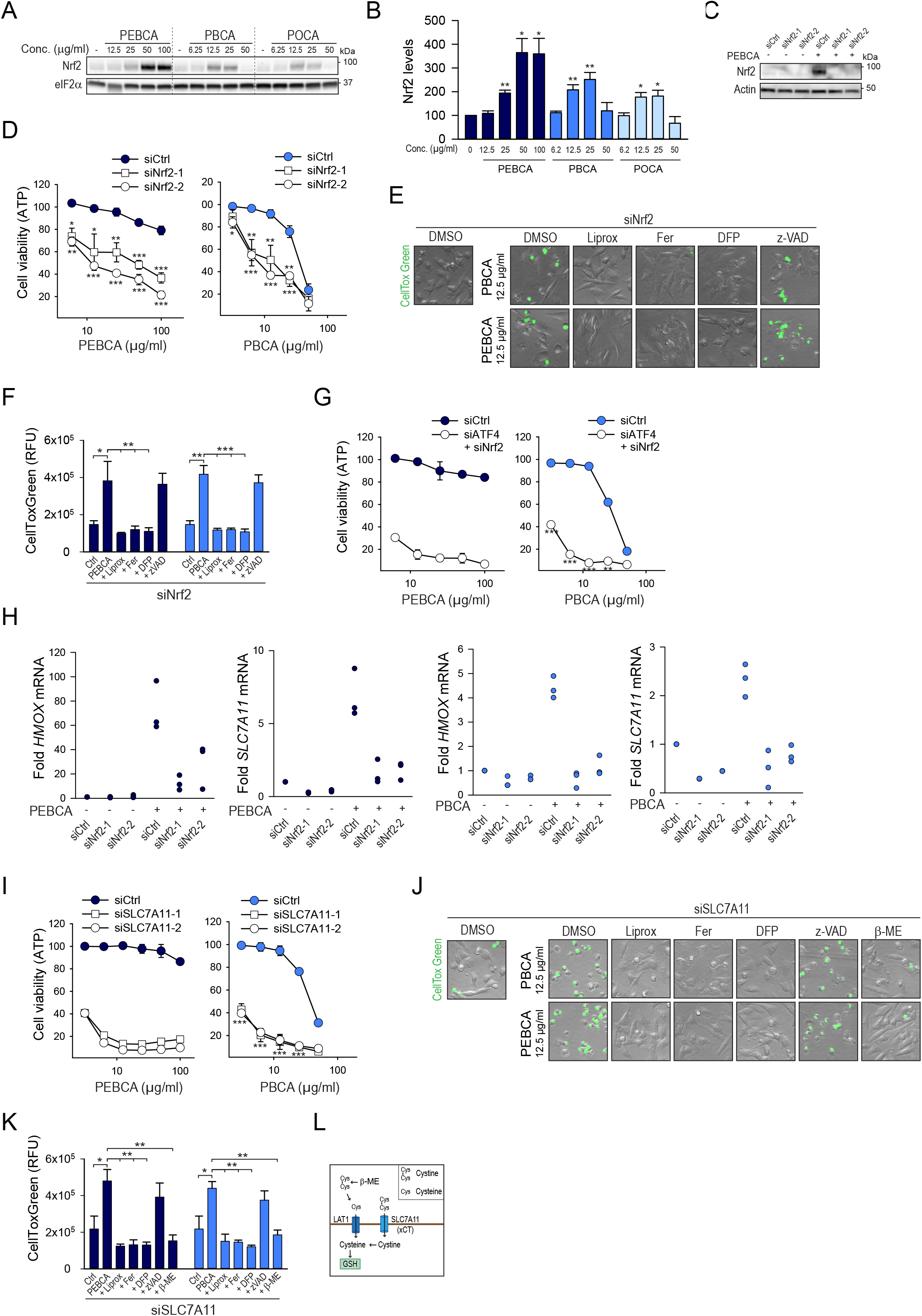
PEBCA - and PBCA-induced ferroptotic cell death depends on ATF4 and Nrf2. **A**. Cells were treated with the indicated concentrations of PEBCA and PBCA for 4 hours and cell lysates were prepared for immunoblotting. The relative levels of Nrf2 were normalized to eIF2α (**B**). **C**. MDA-MB-231 cells were transfected with a non-targeting siRNA (siCtrl) or two independent siRNAs targeting Nrf2. 48 hours later the cells were stimulated with PEBCA particles (50 μg/ml, 6 hours) to demonstrate efficient knockdown of Nrf2. **D**. Cell viability assessed as ATP levels of MDA-MB-231 cells transfected with a non-targeting control siRNA or by two independent siRNAs targeting Nrf2 and 48 hours later treated with PEBCA or PBCA particles for 24 hours. 2-fold dilutions of particles were added, from 100 to 6.25 μg/ml of PEBCA, or from 50 to 3.12 μg/ml for PBCA. Nrf2 depletion in itself reduced cell viability by less than 20%. The data were normalized to the respective non-particle-treated control for each siRNA. **E-F**. Cytotoxicity assessed by CellTox Green staining of MDA-MB-231 cells depleted for Nrf2 and subsequently treated with PBCA or PEBCA particles (12.5 μg/ml, 24 hours) in the presence or absence of liproxstatin (Liprox, 1 μM), ferrostatin (Fer, 2 μM), deferiprone (DFP, 100 μM), or z-VAD (20 μM). **G**. Cell viability assessed as ATP levels of MDA-MB-231 cells depleted for both Nrf2 and ATF4 and 48 hours later treated with PEBCA or PBCA particles for 24 hours. 2-fold dilutions of particles were added, from 100 to 6.25 μg/ml of PEBCA, or from 50 to 3.12 μg/ml for PBCA. The double depletion itself reduced cell viability by 20-25%. The data were normalized to the respective non-particle-treated control. **H**. Transcription of *HMOX* or *SLC7A11* in cells transfected with a non-targeting control siRNA (siCtrl) or by two independent siRNAs targeting Nrf2 and 48 hours later treated with PEBCA (50 μg/ml) or PBCA (12.5 μg/ml) for 4 hours. **I**. Cell viability assessed as ATP levels of MDA-MB-231 cells transfected with a non-targeting control siRNA (siCtrl) or by two independent siRNAs targeting SLC7A11 and 48 hours later treated with PEBCA or PBCA particles for 24 hours. 2-fold dilutions of particles were added, from 100 to 3.12 μg/ml of PEBCA, or from 50 to 3.12 μg/ml for PBCA. **J-K**. Cytotoxicity assessed by CellTox Green staining of MDA-MB-231 cells depleted for SLC7A11 and 48 hours later treated with PBCA or PEBCA particles (12.5 μg/ml, 24 hours) in the presence or absence of liproxstatin (Liprox, 1 μM), ferrostatin (Fer, 2 μM), deferiprone (DFP, 100 μM), z-VAD (20 μM), or β- mercaptoethanol (β-ME, 50 μM). **L**. Schematic overview of cellular import of the dimer cystine and the monomer cysteine. All graphs show mean values ± SEM quantified from at least three independent experiments. The asterisks denote the statistical significances compared to untreated control, unless otherwise indicated. *, p< 0.05; **, p<0.01; ***, p< 0.001.

To investigate whether Nrf2 exerts a protective role in PEBCA/PBCA cytotoxicity, the transcription factor was depleted by siRNA (Figure 6C). Knockdown of Nrf2 rendered the cells highly sensitive to both PEBCA and PBCA particles (Figure 6D), except at high concentrations of PBCA. Moreover, in Nrf2-depleted cells low concentrations of PEBCA and PBCA particles, which were non-cytotoxic in control transfected cells, potently increased cell death in a manner fully reversible by the ferroptosis inhibitors ferrostatin, liproxstatin and DFP, but not z-VAD, implicating induction of ferroptotic cell death upon Nrf2-depletion (Figure 6E-F). This demonstrates that Nrf2 protects the cells against particle-induced ferroptosis. Combined depletion of Nrf2 and ATF4 dramatically sensitized the cells towards both PEBCA and PBCA particles (Figure 6G). Nrf2 has been shown to induce *ATF4* transcription during oxidative stress [28, 29]. In line with the significant activation of Nrf2, PEBCA particles induced ATF4 transcription in an Nrf2-dependent manner (Supplementary Figure 5B), in this way connecting these stress pathways.

Several of the transcriptional targets of Nrf2 are relevant in protection against ferroptosis, such as heme oxygenase 1 (*HMOX1*), ferritin heavy chain (*FTH1*) involved in iron storage, the cystine transporter (*SLC7A11*), or the rate-limiting enzyme in the GSH biosynthesis pathway, glutamate-cysteine ligase (*GCLM*) 14]. Transcription of several of these targets was potently induced by both PEBCA and PBCA in an Nrf2-dependent manner (Figure 6H, Supplementary Figure 5C). To directly explore the significance of the cystine transporter in PBCA/PEBCA-cytotoxicity, MDA-MB-231 cells were depleted for SLC7A11 by siRNA (Supplementary Figure 5D). This strongly sensitized the cells towards PEBCA and PBCA particles (Figure 6I), in a manner similar to double knockdown of Nrf2 and ATF4 (Figure 6G). In cells depleted for SLC7A11, low concentrations of PBCA/PEBCA particles potently induced cell death, in a manner fully reversed by ferroptosis inhibitors, but not z-VAD (Figure 6J-K). To confirm the importance of cystine under these conditions, the cells were treated with β-mercaptoethanol, which reduces extracellular cystine to cysteine that can be imported through the LAT1 transporter (Figure 6L), thus circumventing the need for the xCT transporter. Treatment with β-mercaptoethanol fully reversed the PBCA/PEBCA - induced cell death (Figure 6J-K). Taken together, PEBCA and PBCA particles affect the cellular GSH levels, and compensatory cellular stress responses are induced via activation of ATF4 or Nrf2 to regain homeostasis and prevent cell death.

### PBCA particles induce a transient amino-acid starvation response

From the above data, the PBCA particles more significantly reduced cellular antioxidant capacity than PEBCA particles, and could induce ferroptosis in full medium, whereas PEBCA particles only induced ferroptosis under cystine starvation conditions. In HepG2 cells it was recently shown that a combination of GSH depletion *and* cystine starvation was required for ferroptosis induction, whereas one of the treatments alone was insufficient [30]. Thus, it was of interest to investigate whether PBCA particles affect the cystine availability in addition to depleting GSH. Remarkably, PBCA particles transiently activated the two major amino acid sensing pathways [31, 32], as revealed by activation of GCN2 and suppression of mTORC1 signaling. The particles induced dephosphorylation of the mTORC1 substrate S6K after 15-30 minutes of addition and a transient phosphorylation of GCN2 after 15 minutes (Figure 7A-C). mTORC1 and GCN2 not only sense cellular amino acid levels, but are also the main regulators of cellular protein biosynthesis [33, 34]. Accordingly, in parallel with the PBCA - induced dephosphorylation of S6K, protein synthesis was reduced during the first hour of incubation, before gradual recovery (Figure 7D). As cystine is consumed both in general protein synthesis and in GSH production, a reduction in general protein synthesis would increase the cellular availability of cystine. In line with this, cycloheximide, a well-known inhibitor of protein synthesis, totally abolished the cellular stress responses induced by low concentrations of PBCA under cystine starvation, including the accumulation of ubiquitinated proteins, ATF4, and Nrf2, the phosphorylation of eIF2α, and the dephosphorylation of S6K (Figure 7E). Thus, the transient PBCA-induced amino acid starvation response likely delays particle-induced cytotoxic, oxidative stress effects.

**Figure 7.**
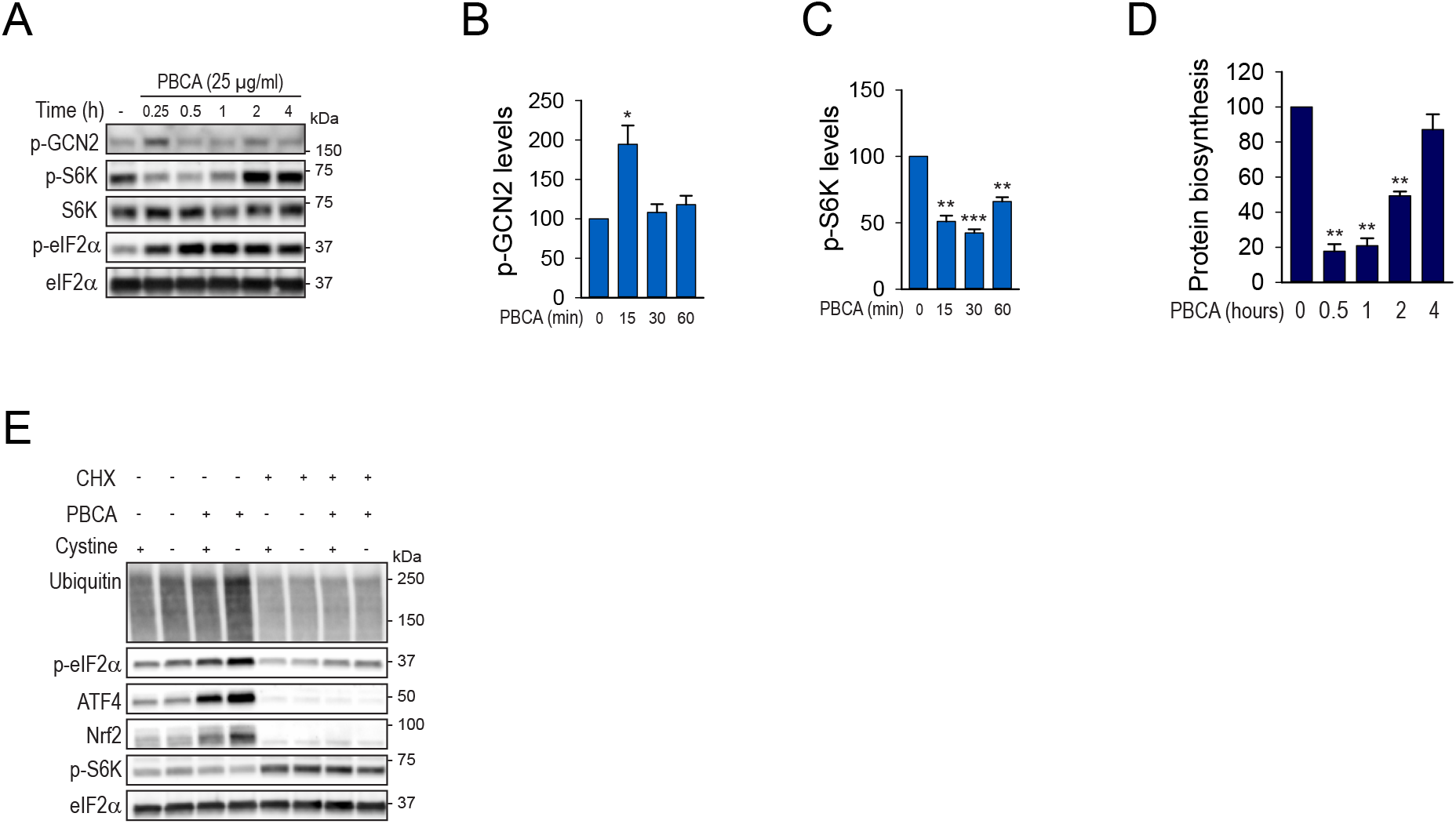
The PBCA particles induce a transient amino acid starvation response. **A-C**. MDA-MB-231 cells were treated with PBCA particles (25 μg/ml) for the indicated times and cell lysates were prepared for immunoblotting. The blots were probed with the indicated antibodies. Relative expression of phospho-GCN2 (B) and phospho-S6K (C) normalized to total eIF2α. **D**. Relative protein synthesis measured by incorporation of [^3^H]leucine for the final 20 minutes in MDA-MB-231 cells treated with PBCA particles for the indicated times. **E**. MDA-MB-231 cells were treated with PBCA particles (12.5 μg/ml) for 4 hours in full - or cystine-free medium in the absence or presence of the protein synthesis inhibitor cycloheximide (CHX, 10 μg/ml) and cell lysates were prepared for immunoblotting. The blots were probed with the indicated antibodies.

## Discussion

Here we have demonstrated that subtle differences in the alkyl side chain of PACA particles have significant impact on particle-induced cellular stress responses and the mode of cell death in cancer cells. Our data may contribute to the choice of PACA monomer as drug carrier in nanomedicine and may help explain the cytotoxicity of these NPs.

The highly similar PACA particles induced surprisingly variable cellular stress responses and cytotoxicity. From our data, it seems unlikely that the three particles merely affect the cells via the same mechanism with different potency. Rather, it seems that each particle induces specific effects during the NP-cell interaction. Moreover, the net effect on cell viability of each particle seems to be determined by the balance between particle-induced pro-death and pro-survival responses. Both the POCA particles and high concentrations of PEBCA particles clearly induced ER stress, as revealed by phosphorylation of PERK, and yet the particle-induced cytotoxicity was very different. Thus, although treatment with both PEBCA and POCA particles affect ER homeostasis, the POCA particles may exert a more acute and severe effect, which renders the cells less able to activate pro-survival pathways to restore homeostasis, and as a result, apoptosis is rapidly induced. In contrast, the PBCA particles did not induce ER stress and PERK activation, but still induced rapid and potent phosphorylation of eIF2α and accumulation of ATF4. This activation was mediated by the ISR, as the accumulation of ATF4 was blocked by the ISR inhibitor ISRIB or simultaneous depletion of the four ISR kinases. We were unable to identify a specific kinase responsible for the PBCA-induced ATF4 accumulation, however, this may be explained by the significant redundancy among ISR kinases [35–37]. As both GCN2, HRI, and PKR have been implicated in detection of oxidative stress [19], and the PBCA-induced ATF4-induction was abolished by both GSH and the antioxidant NAC, it is likely that PBCA-induced oxidative stress contributes to ISR activation. Activation of ATF4 via the ISR exerted a pro-survival role upon treatment with low PBCA concentrations. This may be caused by increased ATF4-mediated transcription of components involved in antioxidant responses, such as the cystine transporter SLC7A11. Furthermore, the PBCA-induced ISR could potentially contribute to cell survival by downregulation of protein synthesis via phosphorylation of eIF2α. Inhibition of protein synthesis would both increase the availability of cysteine for GSH production and reduce the workload for the ER protein folding machinery, thus potentially, limiting both oxidative stress and ER stress. The protein synthesis inhibitor cycloheximide prevented PBCA-induced accumulation of ubiquitinated proteins and activation of the ISR and Nrf2, linking amino acid availability, most likely cystine, to initiation of stress responses. Cycloheximide also abolished the PBCA-induced amino acid starvation response, in line with data from HepG2 cells starved for cystine [30]. Cysteine availability for GSH production may also depend on the extent of amino acid liberation through protein degradation pathways, such as proteasomal degradation. Along these lines, inhibition of proteasomal degradation by bortezomib, a promising anti-cancer drug, induced cell death via limiting the levels of intracellular cystine, and thus GSH, in myeloma cells [38]. Taken together, activation of the ISR constitutes a prosurvival response at low PBCA concentrations. Notably, although the PEBCA-induced ATF4 accumulation is likely caused by the upstream PERK activation, a contribution from the other ISR kinases, for instance by detection of oxidative stress, cannot be excluded.

Our data point to alterations in cellular redox capacity as a key mechanism in the PBCA and PEBCA cytotoxicity. PEBCA-treated cells were able to mount an antioxidant response, such as activation of Nrf2, to a higher degree than PBCA-treated cells. This may contribute to the lower PEBCA cytotoxicity. Nrf2 depletion rendered the cells highly sensitive to both PBCA and PEBCA, possibly by reduced transcription of *SLC7A11*, as depletion of SLC7A11 induced comparable cell death as double knockdown of ATF4 and Nrf2. Increased expression of SLC7A11 is associated with drug resistance and poor survival in several cancer types [39]. Moreover, a subset of triple-negative breast tumors and cell lines, including MDA-MB-231, was shown to require expression of SLC7A11 for growth [40]. In accordance with this, cysteine seemed to play an important role in PBCA/PEBCA-induced cytotoxicity. Cystine starvation reduced cellular GSH levels, increased lipid peroxidation, and strongly potentiated the cytotoxicity of both PBCA and PEBCA particles. It is unlikely that PBCA cytotoxicity is solely caused by inhibition of cystine import, as cystine starvation alone was much less cytotoxic than treatment with PBCA particles. An alternative scenario is that the particles induce concomitant GSH depletion *and* reduced cystine availability, as shown to be required for ferroptosis induction in HepG2 cells [30]. Supporting this, the PBCA particles induced a transient amino acid starvation response, as revealed by inactivation of the mTORC1 substrate S6K and activation of GCN2.

Intriguingly, when the redox capacity of MDA-MB-231 cells was reduced either by cystine starvation or by depletion of Nrf2 or SLC7A11, treatment with PBCA or PEBCA did not induce apoptosis, but rather activated the redox-mediated, iron-dependent cell death mechanism known as ferroptosis. The PBCA particles were able to induce ferroptosis also in cells grown in full medium. These findings are of great interest in drug-delivery, as cellular stress pathways that are induced as a response to the nanocarrier itself may be beneficial if these stress responses sensitizes the target cells. Many cancer types have been shown to be sensitive to ferroptosis induction [41, 42]. Ferroptosis-sensitive cells are characterized by low antioxidant gene expression, including Nrf2 target genes, and decreased levels of GSH and NADPH [15], are highly vulnerable to GSH depletion, and depend on constant cystine import via the Xc - transporter [13]. It was recently demonstrated that inhibition of glutathione peroxidase 4, which is responsible for reduction of peroxidated lipids, leads to ferroptotic cell death of drug-tolerant cancer stem cells in a wide range of cancers [15]. Thus, ferroptosis-inducing compounds have emerged as a novel treatment option for multidrug-resistant cancer stem cells. Ferroptosis-inducing drugs have been shown to cooperate with chemotherapeutics such as cisplatin [43], doxorubicin [44], and gemcitabine [45]. Interestingly, PEBCA particles loaded with doxorubicin are currently in clinical trials for hepatocellular carcinoma (ClinicalTrials.gov NCT01655693), and since PACA particles accumulate in the liver [46], they may influence the redox status of the tumor cells and contribute to cell killing by ferroptosis. Our data also suggests that exchanging the PEBCA monomer with PBCA could potentially prove even more efficient in cell killing of drug-resistant cancers. Moreover, induction of a necrotic tumor cell death such as ferroptosis could potentially contribute to higher tumor infiltration of immune cells, further contributing to tumor clearance [47]. Thus, identification of ferroptosis-inducing NPs is of considerable interest. Certain iron oxide-based NPs have been suggested to contribute to ferroptosis-induction via the Fenton reaction, i.e. iron-catalyzed ROS production [48]. It was recently demonstrated that even small silica-based NPs that are not composed of iron, could induce ferroptosis under amino acid starvation conditions [49]. Based on our findings, it is tempting to speculate that this was due to starvation for cystine specifically, although this was not addressed.

How do NPs induce cellular stress responses, and why do the PACAs studied here have different effects? It has been proposed that physical interactions between NPs and the plasma membrane may signal membrane stress to the cell interior [50]. The interaction between NPs and cellular membranes may lead to alterations in membrane fluidity, the composition of micro-domains, or membrane curvature [51]. Such membrane alterations are known to affect the activity of membrane proteins like receptors, enzymes, ion channels, and nutrient transporters [50]. There are no gross differences in physical characteristics of the three PACA particles. They are of similar size, shape and surface charge. However, their alkyl side chains have increasing hydrophobicity in the order PBCA < PEBCA < POCA. Although the particles are PEGylated, the side chain hydrophobicity may dictate the exact composition of the protein corona that are known to surround all NPs incubated in biological fluids or complete cell medium containing serum [52]. It was recently shown that the hydrophilicity of the NP polymer shell dictated the composition of the protein corona generated in blood [53]. Although the total amount of adhered protein was equal, the protein pattern was different. This determined the NP-cell interactions and the nanocarrier’s stealth properties. Not only proteins, but also lipoproteins have been shown to strongly adhere to NPs [54].Thus, the different cellular effects of PACA particles may be caused by the specific corona surrounding each particle type.

Although the degradation of PACA NPs has been shown to be slow *in vitro*[18], the differential NP cytotoxicity may be caused by release of particle-specific degradation products. Degradation of PACA particles releases cyanoacrylic acid and the corresponding alkyl alcohol, i.e. ethyl-butyl alcohol, butanol, or octanol [55]. It has been indicated that the alkyl chain length dictates the PACA particle degradation rate [18, 55]. POCA particles were shown to have the slowest *in vitro* degradation rate, yet they exert the highest cytotoxicity ([11] and data from this study). Thus, it is tempting to speculate that either the degradation rate is faster when the particles are in contact with the cellular surface than in the absence of cells, and/or the liberated degradation products of the three particle types exert highly different cellular cytotoxicities. In line with this, the hydrophobicity of the released alkyl alcohols is different. The highly hydrophobic octanol will most likely insert into the lipid bilayer and affect membrane properties to a much higher extent than the hydrophilic butanol. In support of this, in neural crests octanol induced intracellular calcium release and apoptosis at *micro*molar concentrations, whereas *milli*molar concentrations of butanol were required [56]. Thus, although POCAs may be degraded more slowly, the liberated alcohol is significantly more potent, which may explain the high POCA cytotoxicity. Moreover, as PBCA particles have the highest degradation rate [18, 55], it cannot be excluded that the GSH-dependent cytotoxicity of PBCA particles, may be caused by liberation of cytotoxic degradation products and cellular GSH depletion through the detoxification process.

In conclusion, even minor differences in PACA particle composition led to differential induction of cellular stress, and importantly, also highly different induction of protective cellular responses like the UPR and antioxidant pathways. Moreover, cystine availability was identified as a key determinant of PBCA/PEBCA cytotoxicity. Importantly, PBCA particles were identified as inducers of ferroptosis, which may be exploited in drug-delivery settings. Together this demonstrates the importance of careful examination of the interaction between NPs and human cells for optimal exploitation of NPs in biomedicine, and to predict nanotoxicity.

## Acknowledgements

We are grateful to Tine Raabe for expert technical assistance, and to Ruth Schmid and Einar Sulheim (both SINTEF, Trondheim, Norway) for valuable discussions concerning PACA toxicity. We thank Maja Radulovic and Prof. Harald Stenmark (both Institute for Cancer Research, Oslo University Hospital, Norway) for the HeLa-mCherry-Galectin-3 cell line. The Core Facility for Confocal Microscopy at Oslo University Hospital is acknowledged for providing access to the confocal microscope, and Core Facility for Mass Spectrometry at OUH for mass spectrometric analyses.

